# Long-Term storage of *Hermetia illucens* (Diptera: Stratiomyidae) larvae

**DOI:** 10.64898/2026.10.12.758660

**Authors:** L. Broeckx, N. B. Lemke, L. Frooninckx, S. Van Miert

## Abstract

The production of black soldier fly larvae (BSFL), *Hermetia illucens* (L. 1758) (Diptera: Stratiomyidae), for organic waste conversion and sustainable biomass production is gaining increasing interest. Maintaining sufficient larval stocks is essential for continuous production, yet practical methods for long-term storage remain limited. This study evaluated the effects of temperature (15°C and 27°C), larval density, and substrate coverage on BSFL survival and growth over a 12-week storage period. Storage conditions significantly affected substrate moisture, larval survival, and development. Covering the substrate with hydrated straw pellets and aluminium foil reduced water loss and minimised maintenance requirements. The highest survival rate, approximately 70% after two months of storage, was obtained at 15°C using this combination of hydrated straw pellets and aluminium foil. Following storage, larvae showed high survival and pupal hatching rates under standard rearing conditions, indicating that prolonged storage did not compromise larval fitness. These results demonstrate that storage at 15°C with hydrated straw pellets and an aluminium cover provides a practical, low-maintenance strategy for maintaining reserve BSFL stocks and ensuring a stable supply of larvae for production and research purposes.

## Introduction

Working with the black soldier fly (BSF) *Hermetia illucens* (L. 1758) (Diptera: Stratiomyidae) can be highly labour-intensive, especially when maintaining multiple BSF strains or when a continuous supply of young larvae is needed. Establishing a buffer stock of larvae would therefore be highly advantageous to ensure stable and uninterrupted BSF cultivation during periods of low fertile egg production, as well as during periods of high workload or limited staff capacity. To enable this, methods for the long-term storage of larvae (i.e., for several weeks to months) need to be developed and adopted. In other fields, such as mass rearing for Sterile Insect Technique (SIT), Integrated Pest Management (IPM) or maggot debridement therapy (MDT), research on long term storage has been conducted and implemented for quite some time. However, the same has not been sufficiently developed for BSFL yet, or at least not in a manner that is publicly accessible. Such efforts may have already been undertaken across the BSFL industry, but often these are tightly guarded trade secrets, and as such an open-source method is not present in the academic literature (discussed below) (Alyagor *et al.*, 2021). Here we discuss some of the theory behind each of these methods and introduce the method that has been implemented and adopted in our facility (Insect Pilot Plant).

### Low temperature

Cold storage of insects is an approach that is commonly used (Colinet and Boivin, 2011; Leopold, 2007), and most methods encompass the initiation and maintenance of reduced metabolism via exposure to temperatures near the species’ lower developmental thresholds. Although cryopreservation, i.e., storage in even lower subfreezing temperatures, can be effectively used for long term storage of the roundworm *Caenorhabditis elegans* (Stiernagle *et al.*, 2006); cryopreservation for insects is much more complicated due to their physiological complexity, and requires a suite of lab-grade chemical agents (e.g., NaOH, NaClO3, Isopropyl Alcohol, hexane, polyethylene glycol, trehalose, liquid N2, etc.) to induce vitrification, which would not be viable for the majority of the BSF rearing industry to employ; though we refer the reader to examples where this has been achieved for other mass-reared Diptera including *Drosophila melanogaster* (Mazur *et al.,* 1992; Zhan *et al*., 2021), the New World screwworm fly *Cochliomyia hominivorax* (Leopold *et al.,* 2001), the house fly *Musca domestica* (Wang *et al*., 2001), and others.

Without a chemically induced cryopreservation process, subfreezing temperatures are typically lethal for most insects, including BSF. For instance, a study examining BSF at subfreezing temperatures, but only *ca.* 2 % of eggs, <1 % of larvae, and no pupae survived after 60 min of exposure at -12 °C (Villazana and Alyokhin, 2019), and indeed freezing is one of the methods commonly used to euthanise insects (Bellezza Oddon S *et al.*, 2025). Therefore, cultured insects like BSF theoretically could be stored under sub-optimal conditions above 0 °C, low enough to stop or decrease development and metabolic rate without reducing survival, adult fitness, and biological function (Colinet and Boivin, 2011). In nature, insects have evolved various strategies to cope with seasonal or acute temperature changes, including diapause, which is a programmed interruption in development in response to low temperatures and decreased metabolic activity (Denlinger, 2023). By artificially inducing diapause, BSFL could putatively be preserved for longer periods.

Because BSFL are ectotherms, the speed of their development is directly related to the temperature of their external environment and the degree-hours they accumulate (Ratte, 1985). Thus, for practical storage, it is crucial to identify a minimum temperature that slows their development but still ensures sufficient survival. The lower temperature threshold for BSFL rearing has been reported by one study to be between 16 and 19 °C (Holmes *et al.*, 2016). In another study, the longest larval and pre-pupal developmental times were recorded at 15 °C, taking 65 days and 86 days, respectively (Chia *et al.*, 2018). At this temperature, larvae showed a survival ratio of 87%; however, the authors noted that no hatching of eggs occurred below 15 °C. Moreover, the absolute cold tolerance of BSF at each life stage has been estimated with and without acclimatisation at 16 °C by measuring the supercooling point (SCP), which is the temperature at which the body liquids freeze, ultimately leading to death. Therefore, the SCP is the absolute lowest limit of cold tolerance of a freeze-avoiding insect (Koch *et al.*, 2004). Late instar larvae had a SCP of -7.28 °C, and pupae had a SCP of -13.7 °C, and each were unaffected by acclimatisation at 16 °C (Spranghers *et al.*, 2017).

### Nutrition

In addition to temperature, diet is known to influence BSFL growth and development (Barragan-Fonseca *et al.*, 2019; Cammack and Tomberlin, 2017; Nguyen *et al.*, 2013). For example, larvae reared on cow dung (which is a low nutrient substrate) at 15 °C had a larval phase lasting 238 days, extending this life stage by nearly 7 months compared to larvae grown on spent grain at 30 °C (Shumo *et al.*, 2019). Moreover, the Sustainable Insect Chain (Susinchain) project, reported that neonates (2-day-old larvae) can be stored with a limited amount of food for 5 days at 25 °C (Sustainable Insect Chain, 2023). Moreover, because access to nutrition for BSFL is modulated by larval density and conspecific competition (Jones & Tomberlin, 2020; Yakti *et al.*, 2022, Rollinson *et al*., 2026), this too is an important parameter to consider. Recent work by Ohhara *et al.* showed that immature BSF larvae can survive starvation for several months and resume growth and development immediately after re-feeding, indicating a starvation-induced quiescent state rather than permanent developmental arrest. The authors further demonstrated that starvation triggered marked atrophy of the fat body, which was fully restored after only five days of re-feeding, suggesting that internal energy reserves are mobilised during starvation to maintain survival and allow rapid recovery once food becomes available (Ohhara *et al.*, 2026).

### Hypoxia

Some research investigating storage and transportation of BSF under low oxygen environments has also been done in recent years. BSFL were stored for 10 days in plastic bags, both with and without vacuum sealing, at either room temperature or 15 °C (Sustainable Insect Chain, 2023). The researchers hypothesised that the absence of oxygen would induce a dormant state in the larvae, reducing their metabolic rate and potentially extending their survival time; however, after 10 days, survival of BSFL in all tested conditions was poor. Based on this research the authors formulated a practical advice that when living larvae need to be stored in their final larval stage for 3 to 7 days, cooling to 15 °C is useful, but vacuum packaging is not. However, because BSFL can survive several days without oxygen in vacuum packaging, the project also investigated survival of BSF larvae in water of different temperatures (room temperature, 15 °C and 3 °C), but larval survival after 3 (72 h) to 6 days (144 h) was lower than survival in non-vacuum packaging (Van Der Borght and Vandeweyer, 2022).

Although several factors influencing BSFL survival and development during storage have been investigated separately, a practical method for maintaining larval reserve stocks over extended periods is still lacking. Therefore, this study evaluated the effects of temperature, larval density, and substrate coverage on BSFL survival and development during a 12-week storage period. In addition, the ability of stored larvae to resume normal development and complete their life cycle after storage was assessed. The aim was to identify a practical, low-maintenance storage strategy that can be readily implemented in both research and commercial BSF production facilities.

## Material and methods

### Provenance

*H. illucens* strains are continuously propagated by the Centre of Expertise Sustainable Biomass and Chemistry (Research group at Thomas More University of Applied Sciences, Belgium) in the Insect Pilot Plant (www.insectpilotplant.be) (IPP) at Campus Geel, as described by Broeckx *et al.* (2021). The Thomas More house strain (TM) was used for this experiment. BSF eggs that had been deposited in traps over a time span of no more than 72 h were harvested from the IPP colony and added to each experimental unit (see below).

### Experimental setup

The experimental unit consisted of a plastic container (height 137 mm, diameter 115 mm, volume 1L, AVA Belgium) filled with 200 g of substrate. To prepare this substrate, 100g of chicken feed (Chicken Start Mash, AVEVE, Belgium) was hydrated with 100 mL tap water to reach a total dry matter content of 45% (Broeckx *et al.*, 2021). To test varying factors to slow the development of BSFL, a 3 × 2 × 3 full factorial design was implemented using n = 2 levels of larval density, n = 3 types of substrate coverage, and n = 2 levels of incubation temperature, with all conditions tested in triplicate, resulting in a total of n = 36 experimental units (Table 1). To create the two levels of larval density, 18 plastic weighing dishes [5mL, VWR, Belgium) were either filled with 0.4 g or 0.8 g of BSF eggs, and these were placed on top of the substrate within the experimental unit as described above. The containers were then initially placed in a climate chamber set at 27 °C and 60 % relative humidity (RH) for 1 week to allow larvae to hatch. Daily, during this period the dish containing the eggs was taken out, and the substrate was gently stirred using a tablespoon to prevent the growth of mould. Afterward, the weighing dish was placed back on top of the substrate. After 3 days, the dishes were removed from the containers, and then 7 days after harvesting the eggs, the following substrate-coverage treatments were applied: (i) no cover (control), (ii) an aluminium foil cover (thickness 102µm, Everyday, Belgium), or (iii) hydrated straw pellets (AVEVE, Belgium) also with an aluminium cover (Figure 1). These hydrated straw pellets were prepared at a mass ratio of 1:3 straw to tap water. 300 g of moistened straw pellets were placed on top of the substrate, but beneath the aluminium cover. Lastly, half of the containers were then split between incubation temperatures of 15 °C, or 27 °C (Pharma 600, Weiss Technik, Germany) for the remaining 11 weeks of the storage experiment, with 60% RH maintained for both) incubation temperatures.

**Table 1.** Experimental design of the storage experiment.

| Eggs | Coverage | Temperature |
| --- | --- | --- |
| 0.4 g | Control | 15 °C |
|  |  | 27 °C |
|  | Aluminium cover | 15 °C |
|  |  | 27 °C |
|  | Straw pellets & aluminium cover | 15 °C |
|  |  | 27 °C |
| 0.8 g | Control | 15 °C |
|  |  | 27 °C |
|  | Aluminium cover | 15 °C |
|  |  | 27 °C |
|  | Straw pellets & aluminium cover | 15 °C |
|  |  | 27 °C |

**Figure 1.**
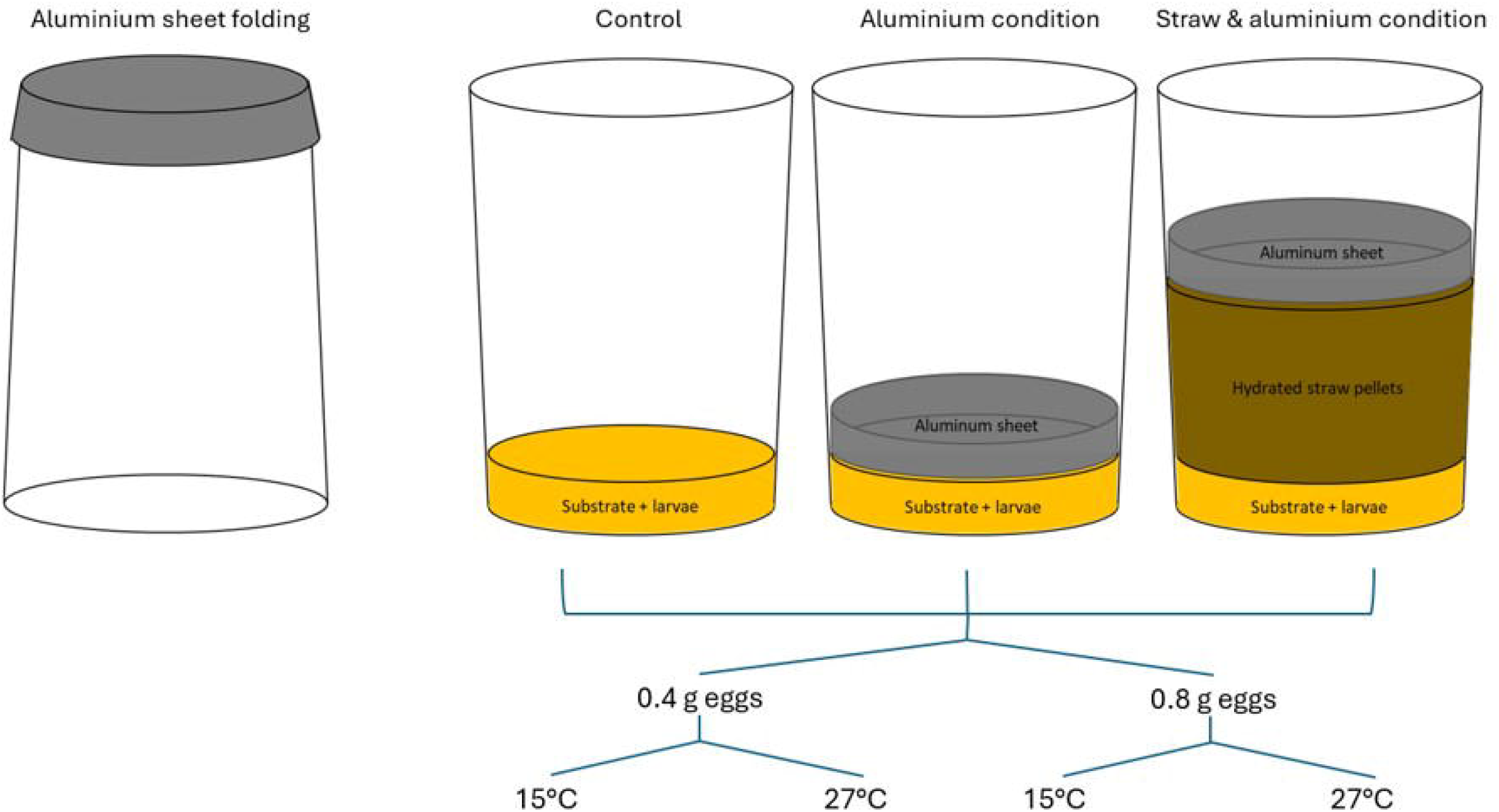
Experimental set-up of the long-term storage experiment. The figure illustrates three experimental coverage treatments: control, aluminium sheet, and straw & aluminium sheet. In each condition, larvae are reared on a substrate, with or without aluminium and hydrated straw pellets, to assess the impact of these variables. For each coverage treatment, two larval densities, obtained by different initial egg masses (0.4 g and 0.8 g), were tested under two different temperatures (15°C and 27°C). The aluminium sheet is folded and placed above the substrate in the aluminium-related treatments.

### Moisture content adjustment during storage

The moisture content of the substrates was monitored and adjusted weekly because the substrate dried out over time. A non-destructive method was used, namely the manual squeeze test (Sharma, 2018). As the name implies, this test is performed by taking a sample of the substrate, squeezing it by hand, and then scoring its compatibility. If the substrate held together as a clump that could later be crumbled without any free water present, the moisture content was considered adequate. When the substrate crumbled immediately after squeezing, this indicated that additional water was required. Whenever this occurred, water was added cautiously until the substrate reached a consistency where it would clump together but could still be crumbled afterward and the total amount of water added was recorded.

### Larval survival and average larval mass during storage

At the start and at weeks 2, 4, 8 and 12 the number of larvae and their weight in each container was assessed by subsampling. To homogenise the mixture to get more accurate subsamples, a miniature mixing apparatus, denoted as the ‘larvae sampling device’, was constructed by employing a rotor operating at 40 RPM and a 6 L bucket mixer drum. The total amount of substrate/frass and larvae was first weighted. Afterwards, the contents were placed into the larvae sampling device and homogenised. 1-3 g of sample was taken, any larvae present were isolated with forceps and then counted and weighed.The total amount of larvae, as well as the average larval mass were then extrapolated from this subsample. Then, to calculate survival, the number of larvae present at the start of the storage experiment was used as the 100 % reference.

### Effect of storage conditions on subsequent larval survival and pupal hatching

To assess the feasibility of the conditions as a long-term storage method, the effect of the storage method on the further survival of larvae and the pupal outcome ratio was investigated. For each of the 12 experiment condition combinations, larvae were pooled from the 3 replicates and 100 were transferred to a container with 20 g of Dry Matter (DM) Chicken Start Mash (CSM), hydrated according to its Water Holding Capacity (WHC) (Frooninckx *et al.*, 2024). After 9 days, larvae were manually counted again to determine the larval survival ratio. After counting the larvae, they were returned to the climate chamber, and 7 days later, 50 prepupae were transferred to empty containers (height 70 mm, diameter 115 mm, volume 0.5L, AVA Belgium) to pupate without substrate as is conventionally done at the IPP. The containers were sealed with a sheet of dry paper towel and a rubber band. Three weeks later, the total number of emerged flies was counted.

### Data analysis

In summary, the responses measured during this experiment were survival, average larval mass, and water addition, recorded at distinct time points over the 12-week trial. For water addition the sum of all additions was calculated to obtain the total amount of water added during the trial. Average and standard deviation from the replicates were calculated for each of these responses. All statistical analyses were conducted using JMP Pro software (version 17.0.0, SAS Institute Inc., Cary, NC).

## Results

The dynamics of substrate mass (including larvae) and the addition of water throughout the storage experiment are presented in **Figure 2**. Across all treatments, a decline in total mass was observed throughout the experiment. The decrease in mass was most pronounced during the first weeks of the experiment. The treatments stored at 27°C had lower substrate masses compared with the treatments stored at 15 °C. Water was periodically added whenever substrates required rehydration to maintain suitable conditions for larval survival. The number of watering periods required were higher at 27 °C compared to 15 °C. Additionally, the type of coverage used (if any) had a large impact on the amount of watering periods. The uncovered treatments consistently required the greatest amount of supplemental water, reaching 89.0 ± 35.9 g by week 4 and 293.2 ± 110.9 g by week 12. In contrast, cumulative water additions remained much lower in the aluminium cover treatments (43.1 ± 15.6 g at week 12) and especially in the straw & aluminium cover treatments (20.1 ± 17.2 g at week 12).

**Figure 2.**
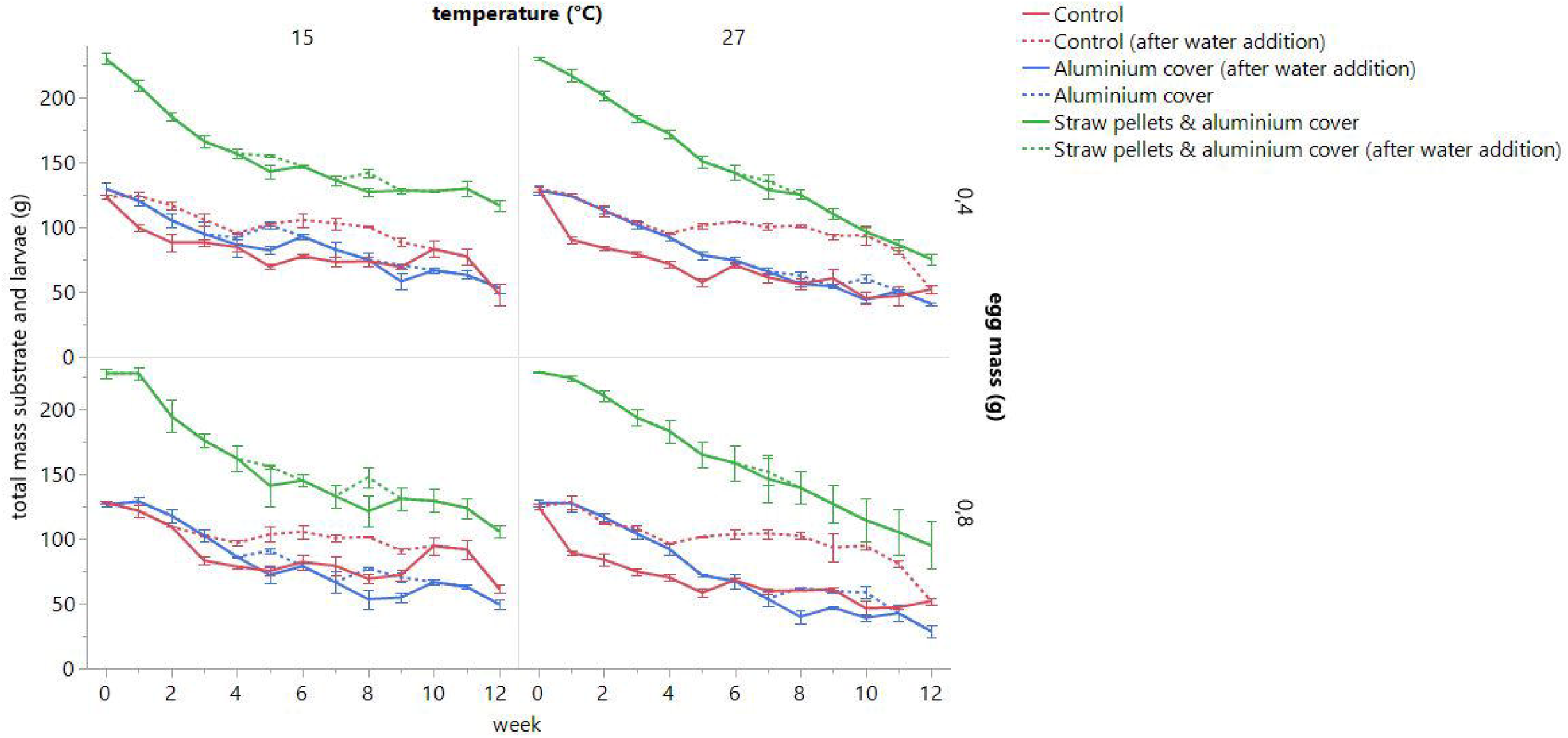
Mass of substrate and larvae (g) over time (weeks) for 3 different substrate-coverage treatments at two temperatures (15 °C and 27 °C) and at two initial egg masses at the start of the experiment (0.4 g and 0.8 g). The four panels compare the effects of the different coverage treatments: control (red), aluminium cover (blue), and straw pellets & aluminium cover (green). Straight lines display the mass of substrate and water before water was added and dotted lines display the mass after adding water. When both a straight and a dotted line are present, it indicates water was added during that week. If only a straight line is present, no water was added that week. Error bars represent the standard deviation of the mean (n = 3).

Larval survival during long-term storage is presented in **Figure 3**. For all conditions, there is a decrease in survival over time. Larval survival tended to be higher at 15°C than at 27°C from the second week of storage onwards. Mean survival at week 2 was 78.9 ± 15.4% for all conditions stored at 15°C compared to 76.6 ± 19.7% for the conditions stored at 27°C. This difference increased over time, with survival rates of 58.3 ± 25.8% versus 50.7 ± 19.4% at week 4, 31.4 ± 19.8% versus 14.0 ± 8.0% at week 8, and 11.5 ± 8.5% versus 3.4 ± 3.2% at week 12 for 15°C and 27°C, respectively. A similar pattern is observed when coverage treatments and initial egg masses were considered separately, with survival consistently higher at 15 °C than at 27 °C from week 8 onwards.

**Figure 3.**
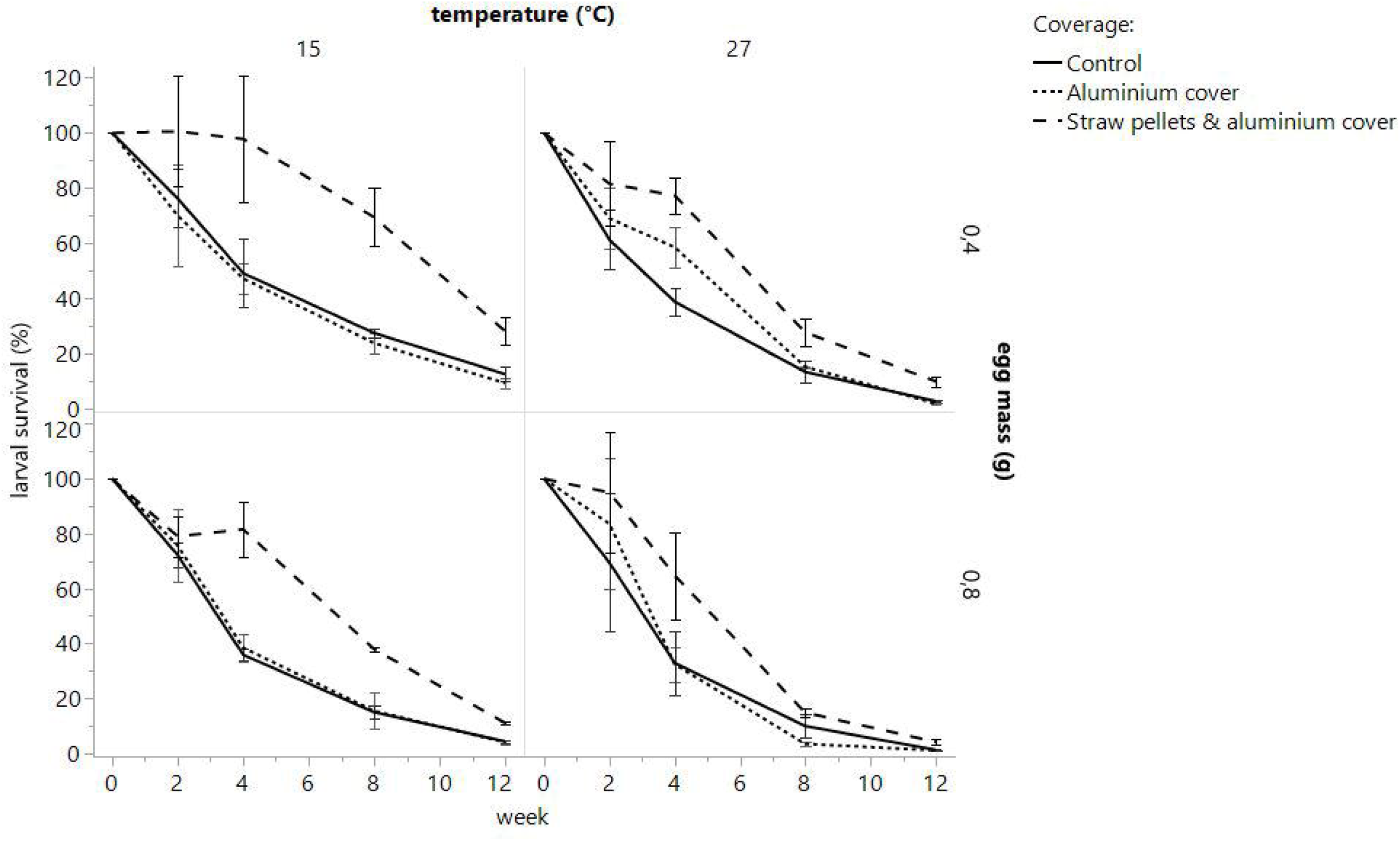
Connecting lines plot of the larval survival (%) over time (week 0-12) for 3 different substrate-coverage treatments at two temperatures (15 °C and 27 °C) and at two initial egg masses at the start of the experiment (0.4 g and 0.8 g). The four panels compare the effects of the different coverage treatments: control (–), aluminium cover (…), and straw pellets & aluminium cover (- - -). Error bars represent the standard deviation of the mean (n = 3).

Larval survival was also influenced by larval densities. At week 4, treatments with a higher larval density had a lower survival (47.64 ± 20.99%) compared to treatments with a lower larval density (61.43 ± 23.08%). At week 8 survival dropped to 16.02 ± 11.34% for the higher larval density treatments and 29.42 ± 19.79% for the lower larval density treatments. By week 12 survival dropped to 4.25 ± 3.47% for the higher larval density treatments and 10.64 ± 9.19% for the lower larval density treatments.

Clear differences for larval survival among coverage treatments were also observed throughout the experiment. When all temperatures and egg masses were analysed together, the treatment consisting of hydrated straw pellets covered with an aluminium sheet consistently resulted in the highest larval survival rates. At week 2, survival reached 89.0 ± 17.4% in the straw & aluminium cover treatment, compared with 74.6 ± 15.9% for the aluminium cover treatment and 69.8 ± 13.9% for the uncovered control. The advantage of the straw & aluminium cover treatment became even more pronounced at week 4, where survival remained at 80.4 ± 18.0%, while the aluminium cover and control treatments declined to 44.1 ± 11.5% and 39.1 ± 9.9%, respectively. Similar trends were observed at week 8, with survival values of 37.4 ± 21.7%, 14.4 ± 8.2%, and 16.3 ± 7.5% for the straw & aluminium cover, aluminium cover, and control treatments, respectively. By week 12, survival remained highest in the straw & aluminium cover treatment (13.2 ± 9.6%), compared with only 4.0 ± 3.5% and 5.1 ± 4.7% in the aluminium cover and control treatments.

The progression of average larval mass during storage is presented in **Figure 4**. Across all treatments, larval mass gradually increased over time. Conditions with fewer initial eggs masses show a higher weight increase, and conditions with a higher temperature also show a higher weight increase. Larval density appeared to have the strongest effect on growth dynamics throughout the experiment. Larvae originating from the lower-density treatments (0.4 g egg mass) consistently achieved a greater average individual mass than those from the higher-density treatments (0.8 g egg mass). At week 2, average larval mass was 7.05 ± 1.56 mg/larva for the 0.4 g treatments compared to 3.19 ± 0.65 mg/larva for the 0.8 g treatments. This difference persisted at week 4 (7.34 ± 1.50 vs. 4.54 ± 1.46 mg/larva), week 8 (10.31 ± 2.17 vs. 6.88 ± 1.87 mg/larva),and week 12 (16.57 ± 4.17 vs. 12.31 ± 1.00 mg/larva for the 0.4 and 0.8 g treatments, respectively). Higher temperatures resulted in higher larval mass during storage. Throughout time this difference increased. At week 2 average larval mass was on average 4.87 ± 2.16 and 5.37 ± 2.43 mg/larva for the treatments at 15°C and 27°C, restively. By week 12 average larval mass was on average 12.68 ± 0.78 and 16.20 ± 4.56 mg/larva for the treatments at 15°C and 27°C, restively. No clear or consistent differences in larval mass were observed among the different coverage treatments.

**Figure 4.**
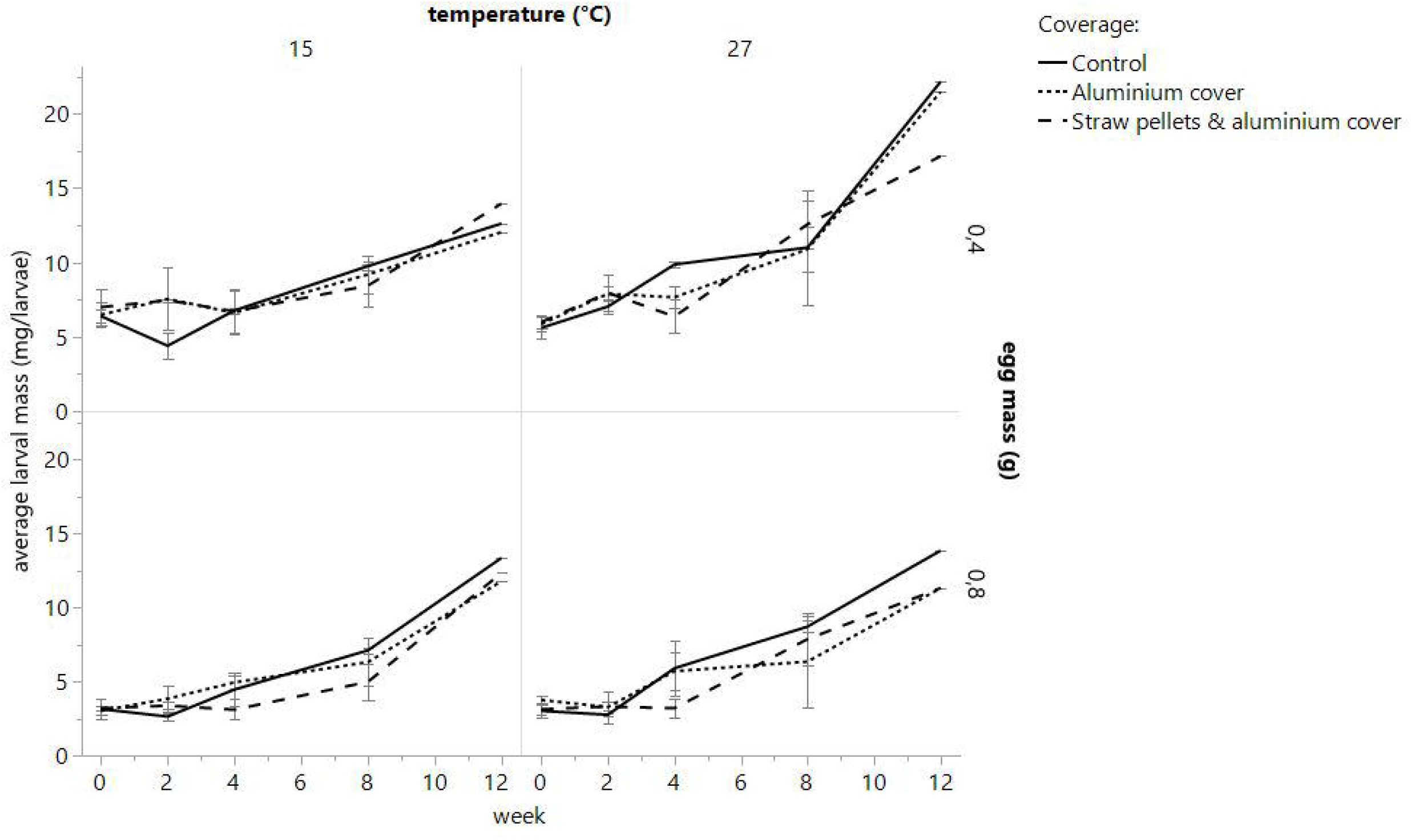
Progression of average larval mass (mg/larvae) over time (week 0-12) for 3 different substrate-coverage treatments at two temperatures (15 °C and 27 °C) and at two initial egg masses at the start of the experiment (0.4 g and 0.8 g). The four panels compare the effects of the different coverage treatments: control (–), aluminium cover (…), and straw pellets & aluminium cover (- - -). Error bars represent the standard deviation of the mean (n = 3).

To determine the effect of the different storage conditions on larval fitness, the survival and pupation ratios of the larvae were assessed after transfer to standard rearing conditions. The results of larval survival and pupal hatching after the storage experiment are shown in **Table 2**. Larvae stored at higher temperatures (27 °C) generally showed higher larval survival and pupal hatching ratios compared to larvae stored at 15 °C. Across all experimental conditions, larval survival ranged from 77% to 99%, while pupal hatching ratios ranged from 84% to 100%.

**Table 2.**
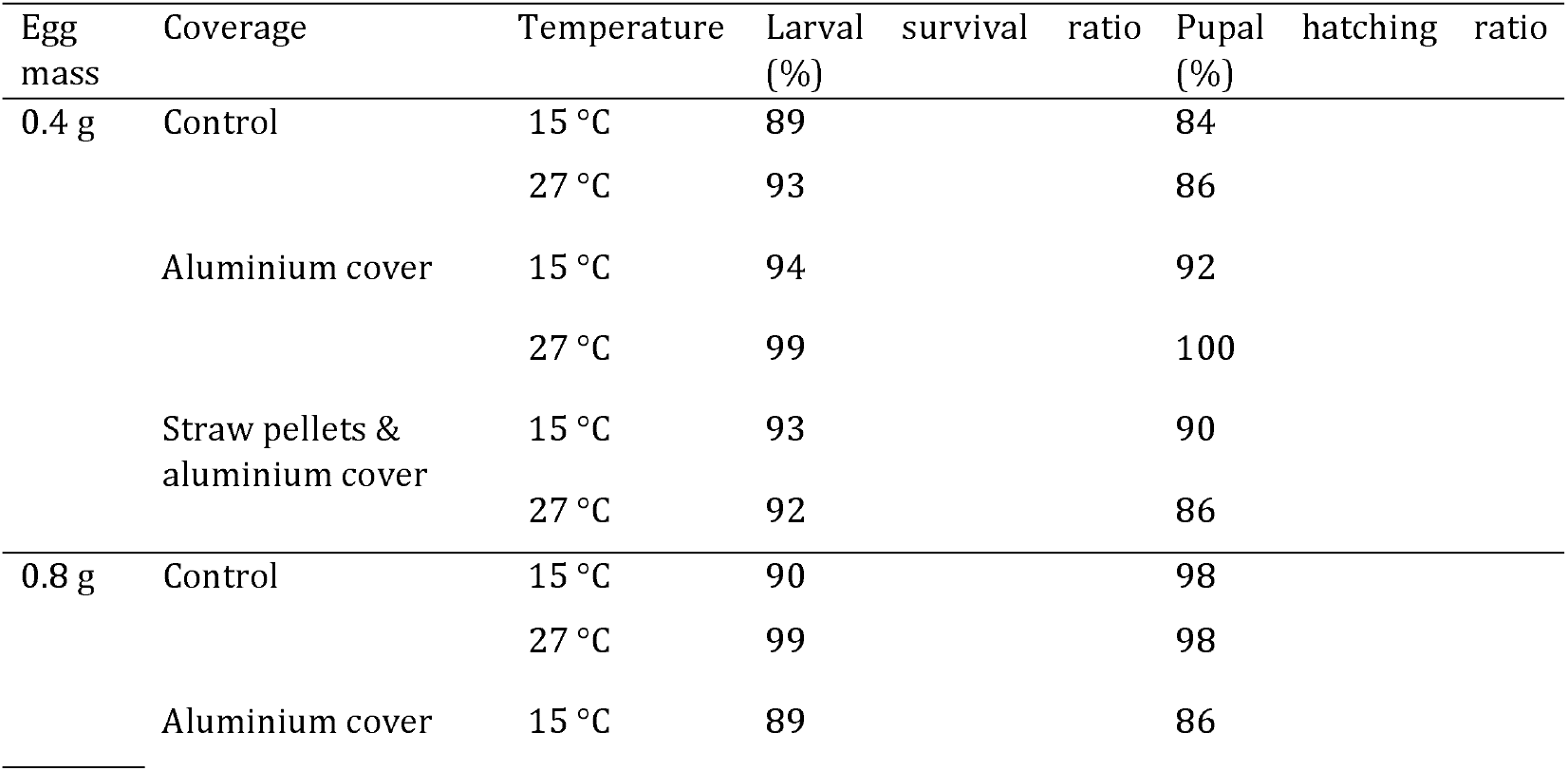
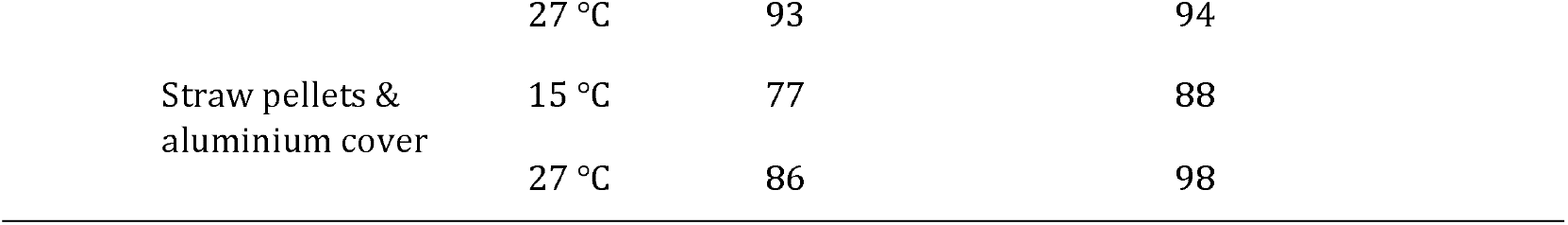
Larval survival and pupal hatching after the storage experiment for each condition.

## Discussion and conclusion

The primary objective of this study was to identify a low-maintenance method for the long-term storage of young BSF larvae. We investigated how storage temperature, larval density (initial egg mass), and coverage influence substrate moisture content, BSFL survival and growth. Moreover, the effects of storage were evaluated by analysing larval survival and pupal hatching under standard rearing conditions after storage for 12 weeks. [for how long?].

### Moisture content adjustment during storage

Substrate moisture content has a pronounced influence on larval growth and survival (Cheng *et al.*, 2017; Frooninckx *et al.*, 2024). Therefore, this parameter was monitored and adjusted throughout the study. Both coverage and temperature affected the amount of water that had to be added during the trial. As expected, the uncovered control required the greatest amount of supplementary watering because the substrate surface was fully exposed to ambient air, increasing evaporative water loss. The addition of an aluminium cover substantially reduced water requirements by limiting the exposed surface area. Furthermore, combining hydrated straw pellets with an aluminium cover resulted in the lowest need for rehydration. The water addition data demonstrate that both covering strategies effectively reduced moisture loss, while the combination of straw pellets and aluminium cover provided the greatest moisture-buffering capacity. The combination of straw pellets and aluminium cover required only two watering events at 15 °C and a single watering event at 27 °C during the entire three-month storage period. From a practical perspective, this treatment is particularly attractive because it significantly reduces maintenance requirements during long-term storage.

During the trial, total substrate mass decreased in all treatments. This reduction most likely resulted from a combination of organic matter consumption by larvae and water loss through evaporation. The most pronounced decrease occurred during the first weeks of storage, potentially because readily available nutrients were still abundant and could be exploited more rapidly during the early stages of larval development. The lower substrate masses observed at 27 °C compared with 15 °C suggest more rapid conversion of substrate dry matter at elevated temperatures. Increased larval and microbial metabolic activity at higher temperatures may have accelerated substrate degradation, resulting in a faster decline in total substrate mass (Bjørge *et al.*, 2018). Consequently, less dry matter remained available to retain water, which may explain differences in watering patterns between temperature treatments.

### Larval weight during storage

For all conditions larval mass increased over time while substrate mass steadily decreased, indicating continued nutrient utilisation. As growth proceeded relatively slowly, this suggests that the storage conditions reduced metabolic activity without completely arresting development (Ohhara *et al.*, 2026).

Temperature had a clear effect on larval growth. Larvae stored at 27 °C generally attained higher individual masses than those stored at 15 °C, indicating that higher temperatures promoted continued development during storage (Bjørge *et al.*, 2018). Larval density also influenced larval growth. Larvae originating from the lower-density treatment consistently achieved higher individual body masses than larvae from the higher-density treatment. The most likely explanation is reduced competition for available nutrients, moisture, and space, allowing each individual greater access to resources (Barragan-Fonseca *et al.*, 2018). Coverage treatment itself had little influence on larval growth, as no consistent differences in larval mass were observed among treatments. This indicates that larval density and temperature were stronger determinants of growth performance than coverage condition. The observation that straw pellets increased substrate dry matter without increasing larval growth further supports the hypothesis that BSFL are unable, or only minimally able, to utilise straw pellets as a nutritional resource, which is consistent with previous studies on highly fibrous substrates (Manurung *et al.*, 2016; Zhang *et al.*, 2023).

Although growth continued throughout storage, larval masses remained below levels typically associated with pupation (Ohhara *et al.*, 2026). Pupae weighing approximately 26 mg have previously been observed in the Insect Pilot Plant. In the present study, this mass was only approached after 12 weeks at 27 °C with an initial egg density of 0.4 g, suggesting that the tested storage conditions effectively delayed development and minimised the risk of premature adult emergence (Davidowitz *et al.*, 2003).

### Larval survival during storage

Larval survival was strongly influenced by both temperature and coverage treatment. Survival was consistently higher at 15 °C than at 27 °C, particularly after prolonged storage [of how long?]. This finding suggests that lower temperatures successfully reduced metabolic activity, allowing larvae to conserve energy reserves and survive longer under storage conditions. In contrast, larvae stored at 27 °C likely experienced higher energy expenditure, resulting in a more rapid decline in survival over time. These findings are consistent with previous storage studies showing that BSFL survival is highest at around 15 °C, while higher temperatures accelerate mortality (Sustainable Insect Chain, 2023).

Larval density also appeared to influence survival, with larvae from the lower-density treatment generally exhibiting higher survival rates than those from the higher-density treatment. Reduced competition for resources may have contributed to this effect, particularly as storage duration increased.

The superior survival observed in the straw with aluminium treatment highlights the importance of moisture preservation during storage. Hydrated straw pellets likely acted as a moisture reservoir while simultaneously restricting evaporation. Combined with the aluminium cover, this created a more stable storage microenvironment that slowed desiccation and supported larval survival. The survival advantage of this treatment became increasingly pronounced with storage duration over 4 weeks, indicating that moisture management becomes particularly important during extended storage periods.

### Effect of storage conditions on larval survival and pupal hatching

The survival ratios of the larvae after storage ranged from 77% to 99%, which is consistent with values reported in previous studies (Broeckx *et al.*, 2021). Similarly, pupal hatching ratios ranged from 84% to 100%, indicating that prolonged storage did not impair successful pupal development (Holmes *et al.*, 2013).

Interestingly, larvae stored at 27 °C generally exhibited higher post-storage survival and pupal hatching compared with larvae stored at 15 °C, despite showing lower survival during storage itself. This suggests that the physiological state of the surviving larvae may differ between storage temperatures. Further investigation is required to determine whether temperature-specific effects on energy reserves, stress physiology, or developmental status contribute to these differences.

### Practical implications

This study demonstrated that young (approximately 1 week old) BSF larvae with individual masses between 4–10 mg can be stored for extended periods of several weeks to months by reducing their metabolism while maintaining adequate substrate moisture. A storage temperature of 15 °C combined with hydrated straw pellets and an aluminium cover substantially improved BSFL survival and formed the basis of the long-term larval storage system currently implemented at the IPP, with minor modifications to improve its practical applicability. During the trials, substrates were regularly mixed to assess moisture content and survival, which likely accelerated dehydration by disturbing the natural moisture gradient within the substrate. Under practical conditions at the IPP, larvae are left undisturbed, resulting in improved moisture retention and successful storage for periods of up to six months. Although lower larval densities resulted in higher survival rates, higher densities remain preferable in practice because they yield a greater absolute number of larvae.

Importantly, larvae surviving prolonged storage successfully resumed normal development, with adult emergence and viable offspring production observed after storage. This confirms that the protocol can be integrated into practical colony management. While the storage system proved effective, further optimisation of storage temperature, substrate composition, and nutrient availability may improve performance, and potential long-term effects on growth and bioconversion efficiency remain to be investigated.

## Acknowledgments

This study was funded by the KU Leuven Industrial Research Fund (IOF) through the IOF C3 project Hermetia Screening Services (C3/24/088). NBL was supported by funds from the US Fulbright Commission to Belgium and Luxembourg under the 2026 US-Belgium Fulbright Scholar Program. Any opinion, findings, and conclusions or recommendations expressed in this material are those of the authors and do not necessarily reflect the views of the US Fulbright Program nor the governments of the United States, Belgium, or Luxembourg.

